# Karyopherin α2 is a maternal effect gene required for early embryonic development and female fertility in mice

**DOI:** 10.1101/2023.06.29.547037

**Authors:** Franziska Rother, Reinhard Depping, Elena Popova, Stefanie Huegel, Ariane Heiler, Enno Hartmann, Michael Bader

**Author notes:** Corresponding author: Franziska Rother. **Summary statement:** We analyze the expression and functional impact of karyopherin α2 in mouse oocytes and early embryos and show that this protein is crucial for the maternal-to-zygotic transition and preimplantation development.

## Abstract

The nuclear transport of proteins plays an important role in mediating the transition from egg to embryo and distinct karyopherins have been implicated in this process. Here, we studied the impact of KPNA2 deficiency on preimplantation embryo development in mice. Loss of KPNA2 results in complete arrest at the 2cell stage and embryos exhibit the inability to activate their embryonic genome as well as a severely disturbed nuclear translocation of Nucleoplasmin 2. Our findings define KPNA2 as a new maternal effect gene.

## INTRODUCTION

Karyopherins are soluble transport factors mediating the nuclear import of proteins. The classical nucleocytoplasmic transport complex involves importin/karyopherin β (KPNB) and one of the six (in mice; in humans and rats one of the seven) known importin/karyopherin α (KPNA) paralogues. The utilization of different KPNA paralogues for nuclear import processes suggests that substrate specificities of KPNAs for different cargoes are existing; on the other hand, dependent on the cell type or tissue, different expression profiles of KPNAs have been found (Kamei et al., 1999; Kohler et al., 1997; Kohler et al., 1999). The differential expression of KPNA paralogues combined with their specificity for distinct substrates can lead to severe effects if one paralogue is depleted (Choo et al., 2016; Liu et al., 2021; Marvaldi et al., 2020; Panayotis et al., 2018; Rother et al., 2011; Sowa et al., 2018; Thiele et al., 2020).

The expression profiles of all known KPNA paralogues have been studied earlier on mRNA level in murine oocytes and the results have shown that only a subset of KPNAs is expressed during oocyte maturation (Mihalas et al., 2015). However, a presence of a distinct KPNA paralogue in the developing oocyte does not automatically stand for its essentiality, as KPNA3 is highly expressed in murine oocytes and its depletion has no effect on oocyte maturation or embryonic development (Rother et al., 2011). However, studies have shown, that the presence or absence of specific proteins in the oocyte has the potential to affect not only the regular development of this highly specialized cell type, but can also impact processes linked to fertilization, zygote formation or start of transcription from the embryonic genome (Mitchell, 2022). Oocyte proteins that have an impact on these processes are called maternal effect proteins, because their expression from the maternal genome is essential for the further development of the embryo independent of the genotype of the embryo. To date, around 70 genes are known to encode for maternal effect proteins and KPNA6/importin α7 has been shown to be one of them (Mitchell, 2022; Rother et al., 2011).

## RESULTS

### Depletion of KPNA2 impacts the ability to give birth to healthy offspring

We analyzed oocytes and zygotes of C57Bl/6 mice to compare the presence of different karyopherin α paralogues. Western blot analysis revealed a strong presence of KPNA2, KPNA3 and KPNA6, while KPNA1 und KPNA4 were not detectable (Fig. 1A). While the absence of KPNA6 in oocytes had been shown to present with a strong phenotype resulting in early embryonic developmental stop, depletion of KPNA3 had been found to be dispensable for embryonic development (Rother et al., 2011) To elucidate the role of KPNA2, which is also strongly expressed in murine oocytes and zygotes, for fertility and embryonic development we created KPNA2 deficient mice (KPNA2 KO; KO). These mice carry a genetrap cassette in intron 3/4 of the KPNA2 gene leading to an aborted transcription of KPNA2 and subsequently formation of a fusion protein containing the importin beta binding domain and β-galactosidase-neomycin fusion protein (Fig. S1A). RT-PCR analysis confirmed the absence of full-length KPNA2 mRNA in different tissues of KO mice, while western blot analysis showed a detectable expression of KPNA2 protein mainly in spleen, thymus, testis, and, to a lesser extent, ovary of wildtype (WT) mice which was absent in KO mice (Fig. S1B,C). KPNA2 KO mice are viable but have a reduced body weight compared to WT littermates. The reduction of body weight was more pronounced in females than in males and continously present during development until adulthood (Fig. S1D).

**Fig. 1.**
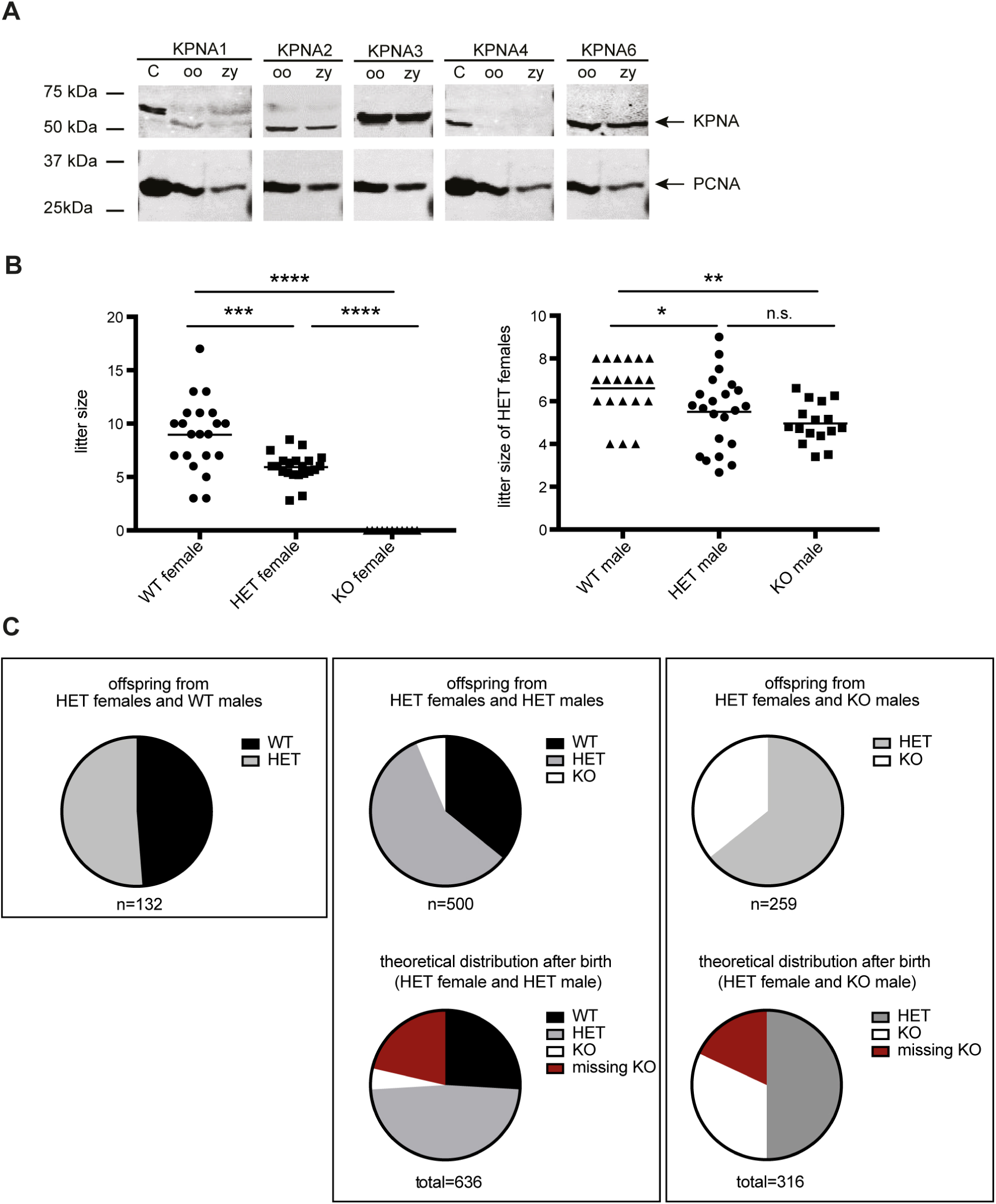
Depletion of KPNA2 impacts the ability to give birth to healthy offspring. (A) Western blot analysis of KPNA paralogue expression in oocytes and zygotes of WT mice. KPNA1 and KPNA4 are absent in oocytes and zygotes, while KPNA2, KPNA3 and KPNA6 can be detected. PCNA served as housekeeping protein. *C* testis protein extract was used as positive control. *Oo* oocytes; *zy* zygotes (B) litter size of KPNA2 deficient females mated with WT males (number of females per group: 11 – 15, number of litters analyzed: n= 11 - 21) and of KPNA2 HET females mated with WT, HET and KO males (number of females per group: 12 – 22, number of litters analyzed: 20 – 102). (C) Genotype distribution of offspring after mating of HET females to WT, HET or KO males; n indicates total number of offspring.

Breeding of KPNA2 KO mice did not result in offspring, while mice heterozygous (HET) for KPNA2 showed a significantly reduced litter size compared with WT (Fig. 1B). Interestingly, the litter size of HET mothers was also dependent on the genotype of males, rising the question, if a partial embryonic lethality could be the reason for the reduced litter size. Analysis of the genotypes of born offspring supported this hypothesis: while breeding of HET females with WT males resulted in offspring according to the mendelian ratios, the breeding of Het females with HET males or with KO males produced a higher ratio of HET offspring than the Mendelian ratios would allow (Fig. 1C). The theoretical recalculation revealed that a part of the KO offspring is obviously missing after birth.

To investigate the fate of the missing KO offspring, we analyzed intrauterine implantation sites of pregnant WT and HET females that had been mated to WT males. We found a slightly lower number of implanted embryos at E11.5-E13.5 in HET females, while there was a tendency to having more empty implantation sites in these mice (Fig. S2A,B). The morphological analysis of implanted embryos showed no statistical significance when comparing the fraction of avital or developmentally retarded embryos in WT and HET females (Fig. S2C). However, genotyping of implanted embryos of HET females revealed that smaller (delayed) embryos had a higher chance of being KPNA2 KO than larger embryos (Fig. S2D). This developmental delay in KPNA2 KO embryos was still present at E17.5 -E20.5. We conclude that, as no major hints for intrauterine embryo loss were present, a part of KPNA2 KO offspring could die during the birth process, while being smaller and maybe weaker than their heterozygous siblings.

### Depletion of KPNA2 leads to a reduced formation of fertilizable oocytes

Next, we investigated the reason for infertility of KPNA2 KO females. After hormonal stimulation, KPNA2 KO mice presented a dramatically reduced number of both MII oocytes and GV-oocytes compared to WT females (Fig. 2A,B). Although we did not observe significant differences in the chromatin status of GV-oocytes between the groups (data not shown), in vitro maturation of GV-oocytes produced an even reduced number of MII oocytes in KPNA2 KO females while HET females were unaffected (Fig. 2C). Additionally, the MII oocytes retrieved after hormonal stimulation frequently showed abnormal spindle formation in the KPNA2 KO group compared to WT and HET (Fig. 2D,E). The poor quality of oocytes from KPNA2 KO females could be verified by a tendency to activate spontaneously (Fig. 2F). Together these data suggest that KPNA2 is necessary for female fertility by impacting the regular development and maturation of oocytes.

**Fig. 2.**
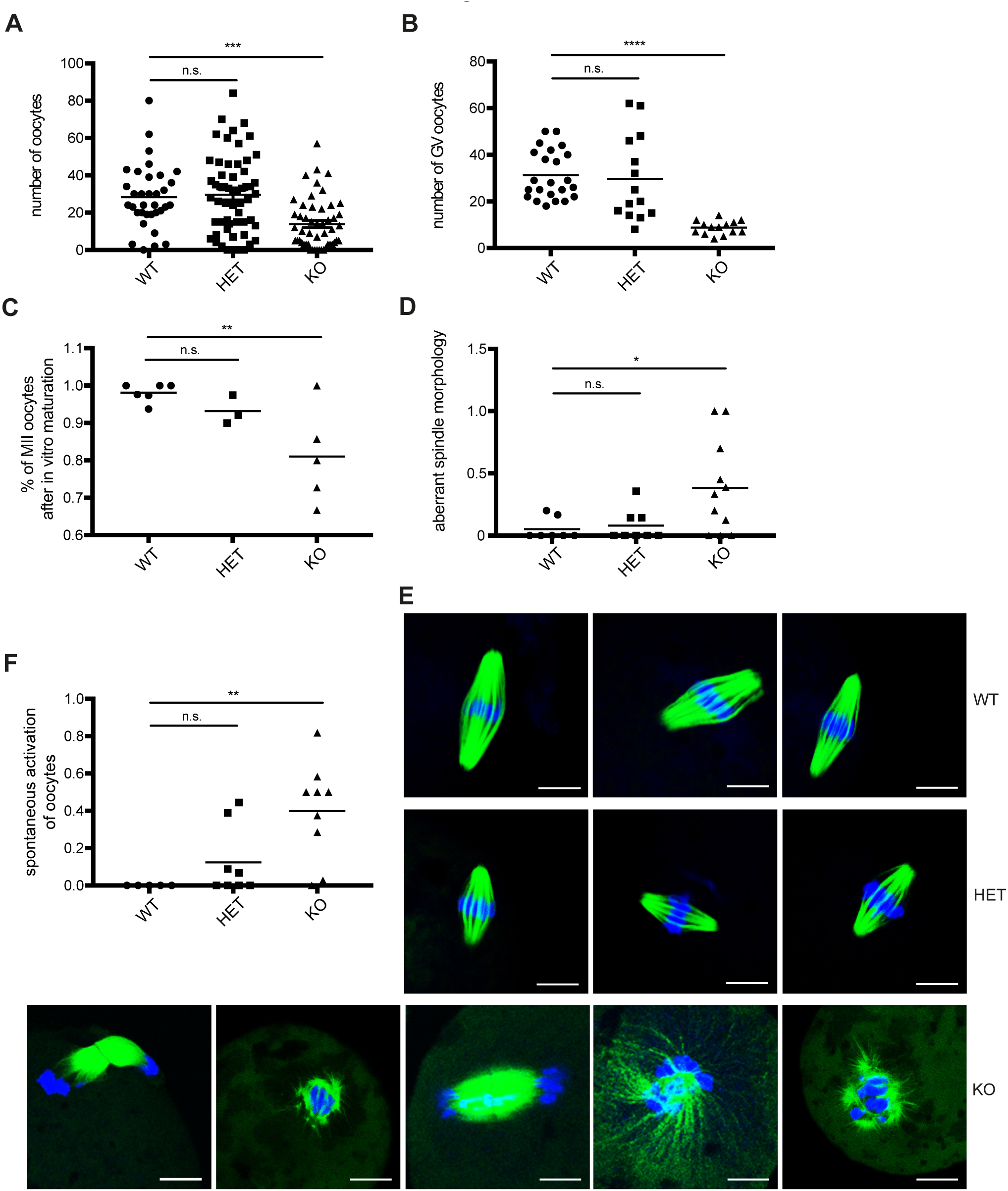
Depletion of KPNA2 leads to reduced formation of fertilizable oocytes and aberrant spindle formation. (A) number of oocytes retrieved after hormonal superovulation of mice (n = 18 - 42 females / group). (B) number of GV oocytes retrieved after hormonal superovulation (n = 14 - 23 females / group). (C) Fraction of MII oocytes retrieved after in vitro maturation of GV oocytes (n = 3 - 6 females / group). (D) Fraction of oocytes with visually abnormal spindle morphology (n = 7 - 11 females / group, number of oocytes analyzed: 74 – 103 / group). (E) Immunofluorescence of MII spindles showing tubulin (green) and DAPI (blue). Scale bar 10 μm. (F) Fraction of oocytes showing spontaneous activation determined by formation of a pronucleus (n = 5 - 9 females / group, number of oocytes analyzed: 101 – 171 / group).

### Maternal depletion of KPNA2 leads to an arrest in early embryonic development

As the above described phenotype does not completely explain the fact that no single offspring could be retrieved from KPNA2 KO females, we investigated the first developmental steps after fertilization of oocytes. Interestingly, the few retrieved oocytes of KPNA2 KO mice showed a normal fertilization rate defined by pronuclear formation and extrusion of the polar body and development into 2cell embryos occurred properly (Fig. 3A,B). However, none of the 2cell embryos was able to undergo a further cell division, concluding that in addition to a subfertility maternal KPNA2 deficiency leads to an early embryonic developmental arrest (Fig. 3C,D). Analysis of the developing zygotes revealed an elevated rate of zygotes with abnormal formation of pronuclei: either one of the two pronuclei was missing or more than two pronuclei could be found (Fig. 3E). However, the developmental stop at the two cell embryo stage occurred not only in embryos with abnormal pronuclear numbers, but in all embryos derived from KPNA2 knockout mothers.

**Fig. 3.**
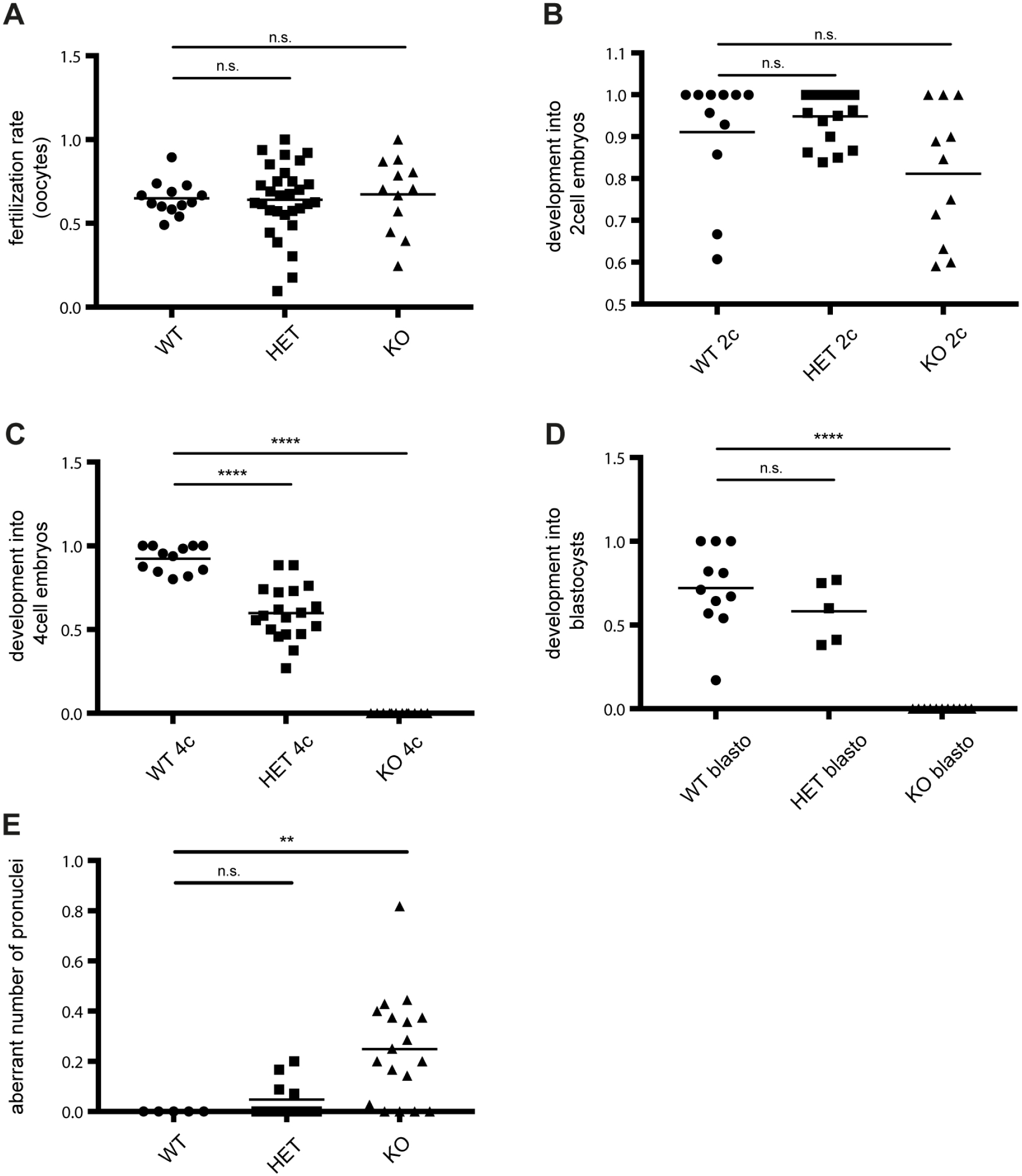
Maternal depletion of KPNA2 results in early embryonic arrest. (A) Normal fertilization rate in oocytes from KPNA2 KO mice. Number of mice: 12 – 30 / group; number of oocytes: 337 – 1020 / group. (B) Fertilized eggs of KPNA2 KO females show regular cleavage into 2cell embryos. Number of mice: 11-17 / group; number of eggs: 147-352 / group. (C, D) Embryos from KPNA2 KO females don’t develop beyond 2cell stage. Number of females: 5 – 19 / group; number of embryos: 113 – 331 / group. (E) Zygotes of KPNA2 KO females frequently display abnormal pronuclear formation resulting in aberrant number of pronuclei. Number of mice: 5 – 18 / group; number of zygotes: 122 – 156 / group.

### KPNA2 protein is translated from stored maternal mRNA in zygotes

To follow the activity of the KPNA2 promoter, which in KPNA2 KO mice regulates the expression of the IBB-β-galactosidase fusion protein, we performed an X-gal staining in oocytes and early mouse embryos (Fig. 4). GV oocytes of heterozygous and KPNA2 KO mice showed robust activity of β-galactosidase concluding that KPNA2 is regularly expressed at this stage. In MII oocytes, no β-galactosidase activity could be found, most likely due to degradation of the artificial fusion protein. However, zygotes of heterozygous und KPNA2 KO females that had been mated to WT males showed a de novo expression of β galactosidase protein. To discriminate at this stage between a new transcription from the embryonic genome vs. translation from stored maternal mRNAs, we mated WT females with KPNA2 KO males. No β-galactosidase expression could be detected in these zygotes, suggesting that β-galactosidase as a representative of KPNA2 is translated from stored maternal mRNAs. Moreover, after mating of heterozygous females with WT males 98% of the retrieved zygotes showed positive β-galactosidase signals further supporting the hypothesis of translation from maternal mRNAs (data not shown). In 2cell embryos the Xgal staining finally revealed the newly expressed KPNA2 equivalent, as 2cell embryos from WT females mated with KPNA2 KO males stained positive. We conclude that KPNA2 expression from the embryonic genome starts at this early stage. Interestingly, the expression from the KPNA2 promoter was found to be much weaker in zygotes derived from KPNA2 KO females compared to zygotes derived from KPNA2 heterozygous females. This finding indicates an impaired embryonic genome activation in embryos derived from KPNA2 KO mice.

**Fig. 4.**
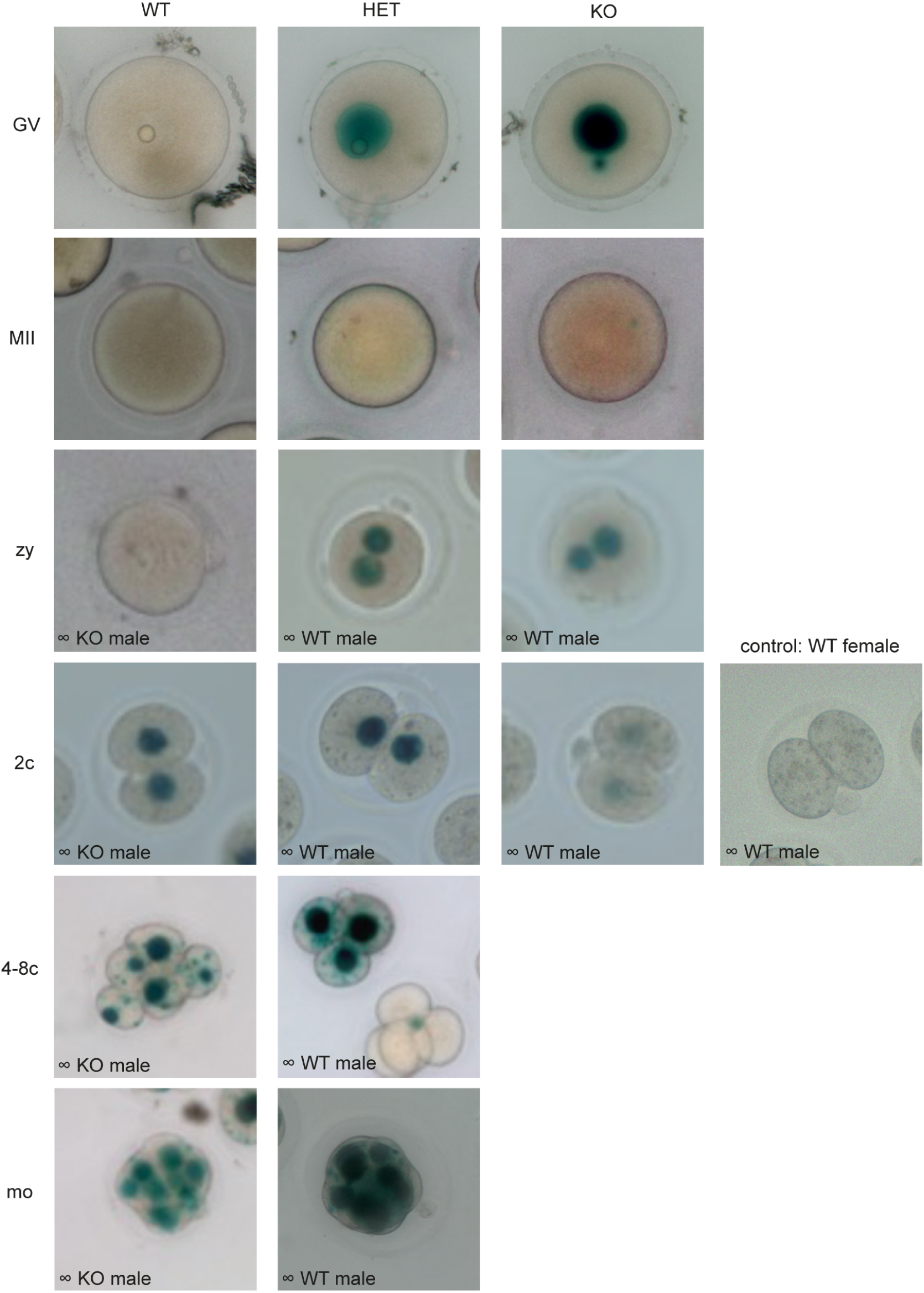
KPNA2 promoter activity during early embryonic development. X-gal staining of early embryos at different stages of development reveals formation of the artificial IBB-β-galactosidase fusion protein. The column labeling (WT; HET; KO respectively) refers to the genotype of the female the oocytes are derived from. Eggs and embryos carrying at least one genetrap allel for KPNA2 have the potential to form the fusion protein. KPNA2 (-equivalent) is formed in GV-oocytes and rapidly dissolves in MII oocytes. In zygotes, the formation of the fusion protein is dependent on stored maternal mRNAs, while starting in 2cell embryos, a de novo formation of the fusion protein can be detected. No 4-8c and morula stage embryos could be retrieved from KO females, as embryos display a 2cell stop. *GV* GV stage oocytes; *MII* MII stage oocytes; *zy* zygotes; *2c* 2cell embryos; *4-8c* 4cell – 8cell embryos; *mo* morula stage embryos. The inserted text in the photos depicts the genotype of males used for breeding.

To exclude a dominant negative effect of the IBB β-galactosidase fusion protein on early embryonic development, we tracked the development of embryos derived from mating of KPNA2 heterozygous females with WT males (Table S1). While 98% of zygotes showed a positive X-gal signal, 49% of 2cell embryos and 55% of 4cell embryos were positive. A dominant negative effect would have led to an enhanced death rate of X-gal positive 2cell embryos, thereby reducing the rate of positive X-gal signal in surviving 4cell embryos. Thus, we can exclude a negative effect of the IBB-β-galactosidase fusion protein on early embryonic development.

### Localization of KPNA2 in embryos is dependent on the developmental stage

To analyze the expression and localization of KPNA2 in WT preimplantation mouse embryos, we performed immunohistochemistry for KPNA2 in embryos at different developmental stages (Fig. 5). In GV oocytes, KPNA2 was moderately expressed and localized in the GV as well as in the ooplasm. In MII oocytes, KPNA2 was redistributed in the ooplasm with no specific localization to the MII spindle, while in zygotes, the protein strongly localized to both pronuclei. After first cleavage into 2cell embryos, KPNA2 expression was found in cytoplasm and nuclei with a particular localization at the nuclear membrane. This strong localization to the nuclear membrane was also seen in later developmental stages until morula. In blastocysts, KPNA2 displayed a dotted signal and distributed in the cytoplasm and nuclei with no specific enrichment. The strong pronuclear localization of KPNA2 in zygotes led to the conclusion for a very specific function of KPNA2 at this developmental stage.

**Fig. 5.**
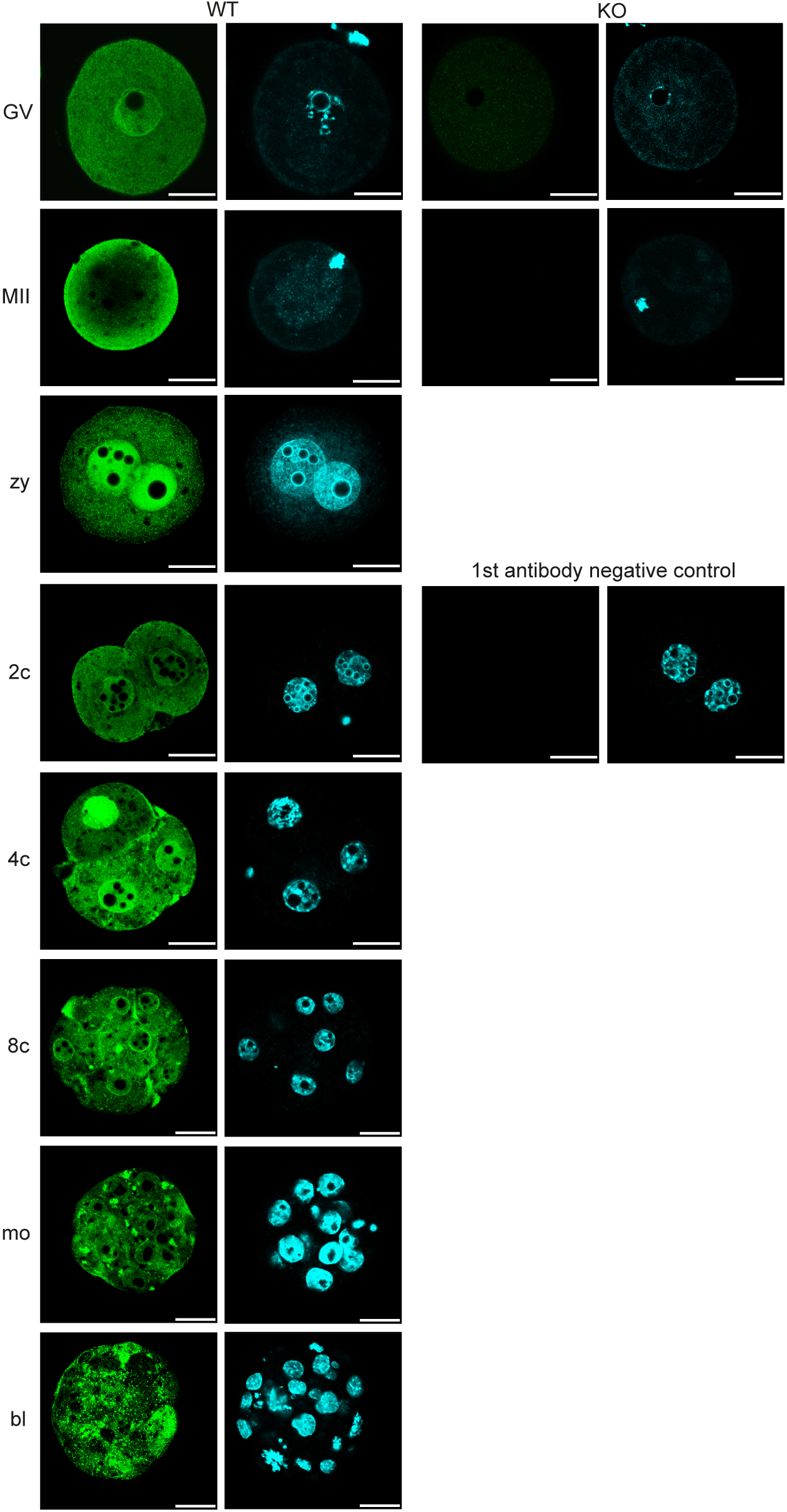
Localization of KPNA2 in developing preimplantation embryos. Immunofluorescence staining of KPNA2 (green) in zygotes reveals a strong signal in both pronuclei. After cleavage into 2cell embryos the spatial distribution of KPNA2 temporally alternates between localization in the cytoplasm, nuclear accumulation and distinct signals at the nuclear membrane. DNA is visualized with DAPI (blue). *GV* GV stage oocytes; *MII* MII stage oocytes; *zy* zygotes; *2c* 2cell embryos; *4c* 4cell embryos*; 8c* 8cell embryos; *mo* morula stage embryos; *bl* blastocysts; scale bar 25μm.

### Absence of KPNA2 impairs the zygotic genome activation (ZGA)

To test, whether the activation of the embryonic genome is affected in embryos derived from KPNA2 KO females, we performed RT-PCR for several genes that are known to be markers of ZGA (Hamatani et al., 2004; Zeng and Schultz, 2005). 2cell embryos derived from KPNA2 KO females displayed a massively reduced expression of the eukaryotic translation initiation factor 1A (EIF-1a) and no expression of KPNA1 and nucleoside diphosphate kinase 2 (Nme2, Fig. 6). In contrast, Kpna3, which has been shown before to be constitutively expressed from oocytes throughout early embryonic stages, was shown to be normally expressed in 2cell embryos of KPNA2 KO females (Rother et al., 2011; Wang et al., 2023). Thus, we conclude that embryos from KPNA2 deficient females cannot activate their genome.

**Fig. 6.**
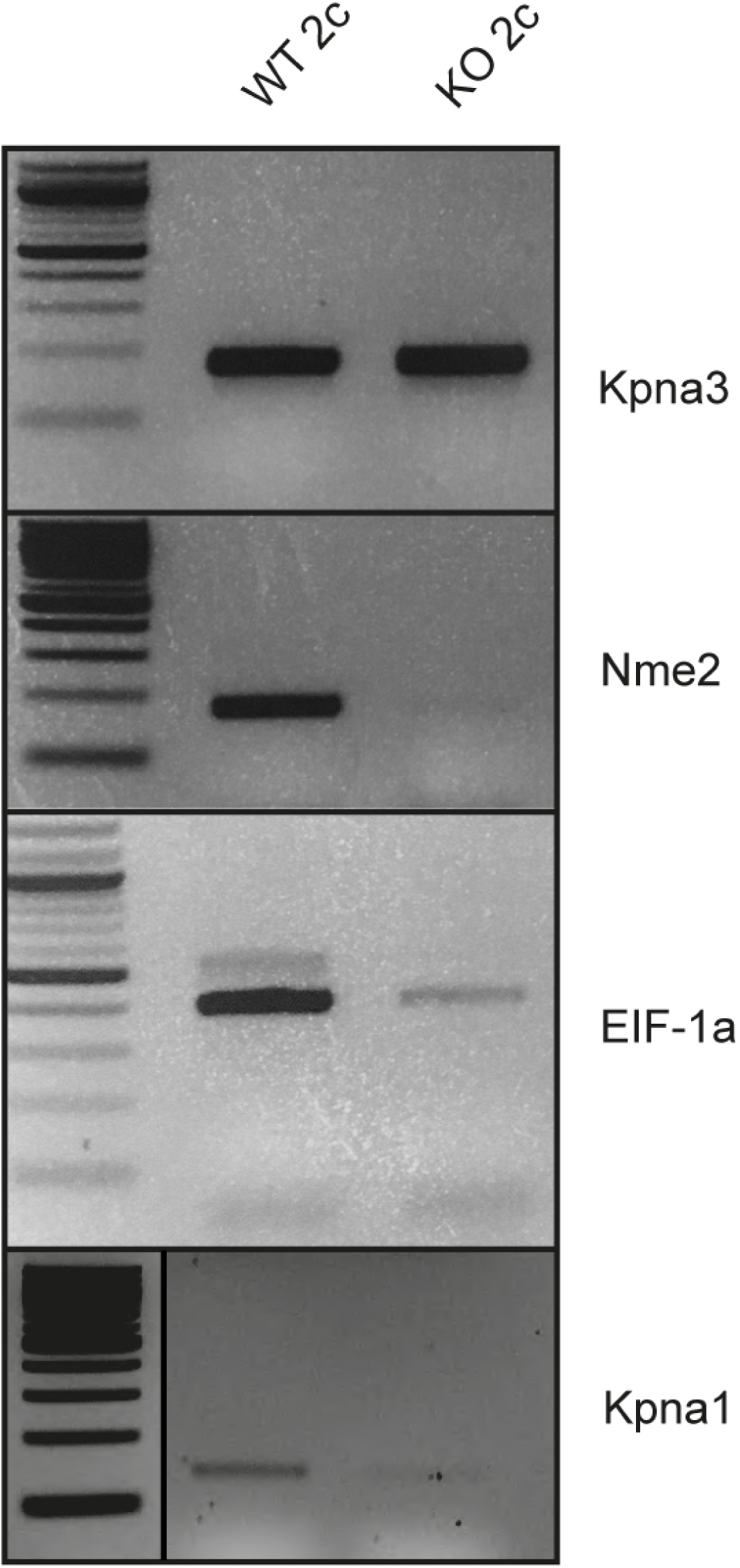
Impaired ZGA in KPNA2 KO embryos. RT-PCR of 2cell embryos shows a reduced or missing expression of the ZGA-dependent genes Nme2, EIF-1a and Kpna1, in KPNA2 KO derived embryos. In contrast, Kpna3 mRNA which is also maternally expressed, is still detectable in KPNA2 KO derived 2cell embryos.

### Regular nuclear expression of the transcription factor Oct4 in KPNA2 KO embryos

We hypothesized that the impaired ZGA could be a result of the aberrant expression or localization of a transcription factor. The transcription factor Oct4 is a key regulator of the first wave of ZGA and its depletion leads to an early developmental arrest in mice (Tan et al., 2013). Previous studies have shown that Oct4 possesses a nuclear localization sequence (NLS) which is essential for its nuclear import and that the transcription factor binds to KPNA2 (Li et al., 2008; Pan et al., 2004). We thus hypothesized that Oct4 localization could be affected in embryos from KPNA2 KO females. However, immunohistochemistry with an antibody against Oct4, detecting both isoforms revealed a regular localization of the transcription factor in the GV, pronuclei and nuclei of WT and KO oocytes and embryos (Fig. 7A). Thus, the absence of maternal KPNA2 did not lead to a defective import of Oct4 in these embryos. To further investigate the reason for an undisturbed Oct4 localization in KPNA2 KO embryos, we performed a binding assay to test, whether other KPNA2 paralogues were able to bind to Oct4. Interestingly, all tested KPNA paralogues and importin β bound to Oct4 (Fig. 7B). The binding of Oct4 and KPNA paralogues was facilitated when importin β was added and reduced by adding RanGTP which proves the specificity of the binding. Thus we concluded, that KPNA2 is not exclusively transporting Oct4 into the nucleus.

**Fig. 7.**
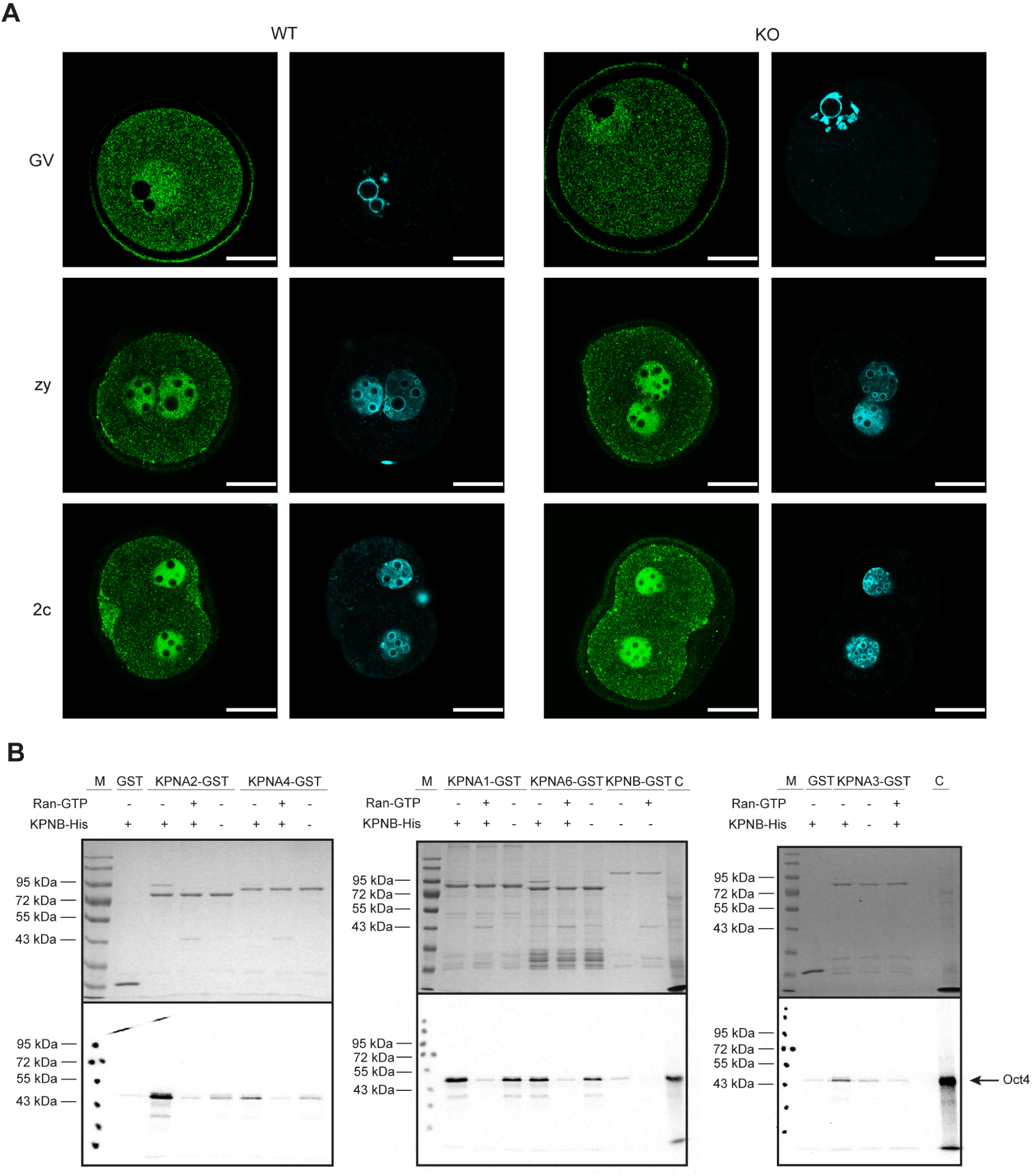
Normal nuclear expression of the transcription factor Oct4 in KPNA2 KO embryos. (A) Immunofluorescence staining of Oct4 reveales a regular localization in germinal vesicles, pronuclei and nuclei (green); DNA is visualized with DAPI (blue). Scale bar 25μm. (B) Oct4 binding assay using KPNA-GST and ^35^S-methionine labelled in vitro transcribed and translated (IVTT) Oct4. The binding assay confirms a specific Importin β dependent binding of all tested KPNA paralogues to Oct4. *M* marker; *C* positive control (IVTT Oct4 10% input).

### Nuclear transport of nucleoplasmin 2 is disturbed in zygotes and 2cell embryos of KPNA2 deficient females

A recent report by Wang et al. had suggested that nuclear import of the protein RSL1D1 is impaired in KPNA2 KO embryos and that this could account for the developmental stop of embryos, especially, since the RSL1D1 KO showed a similar phenotype as the KPNA2 KO (Wang et al., 2023). To understand to what extend RSL1D1 is involved in the 2cell embryonic stop, we analyzed its nuclear localization in zygotes and 2cell embryos. Interestingly, we noticed a high variability in nuclear RSL1D1 signals in WT and KO zygotes with some zygotes showing weaker nuclear enrichment than others (Fig. 8A,C). Further analyses in 2cell embryos showed no clear differences in nuclear localization of RSL1D1 in WT and KO, possibly due to the observed high variability.

**Fig. 8.**
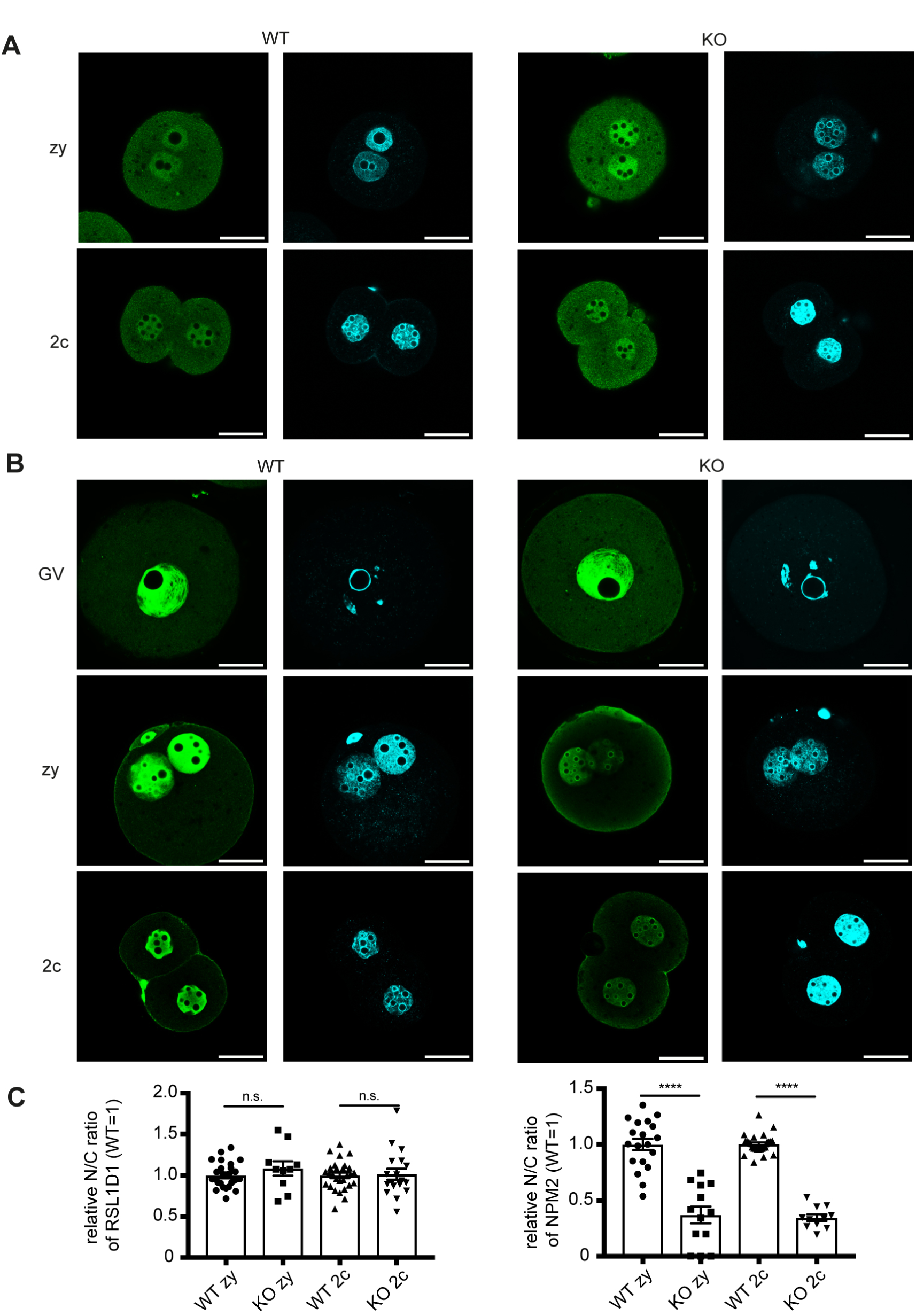
Nuclear localization of the maternal effect proteins RSL1D1 and NPM2 in oocytes and early embryos. (A) Immunofluorescence staining for RSL1D1 (green) and DAPI (blue) in zygotes and 2cell embryos of WT and KPNA2 KO mice. (B) Immunofluorescence staining for NPM2 (green) and DAPI (blue) in GV oocytes, zygotes and 2cell embryos of WT and KPNA2 KO mice. (C) The ratio of nuclear and cytoplasmic signals was quantified. Number of animals quantified for RSL1D1 signal: 2 - 3 / group; number of cells: 10 – 28 / group. Number of animals quantified for NPM2 signal: 3 - 4 / group; number of cells: 11 – 26 / group. Scale bar 25μm.

We hypothesized that multiple factors could be causative for the 2cell stop in KPNA2 KO embryos, as nuclear transport is elementary at that stage and deficiency in other KPNA paralogues have shown to be fatal for embryonic development as well (Rother et al., 2011). In search for additional factors responsible for the developmental stop and disturbed ZGA in KPNA2 KO embryos, we tested the localization of multiple known factors that have been described to act as maternal effect proteins. The transcription factor Sall4 which is known to interact with Oct4, the transcriptional activator Brg1/Smarca4, which modifies the chromatin structure, the germ cell specific protein Stella/Dppa3 known to be involved in the epigenetic chromatin reprogramming in the zygote following fertilization, as well as HDAC1, whose proper function prevents a 2cell developmental stop in mouse embryos were analyzed and none of these factors showed an altered spatial distribution in early embryos which would explain the developmental arrest (Fig. S3). Moreover, the methylation of histone H3 at lysin9 which has been shown to be important for coordinated ZGA did not reveal any differences between WT and KO zygotes (Fig. S3).

In contrast, we found nucleoplasmin 2 (NPM2) to be differentially located in KPNA2 KO embryos. NPM2 is a histone chaperone involved in chromatin reprogramming, especially during fertilization and early development and it is highly abundant in oocytes and zygotes where it has been shown to act as a maternal effect protein in mice, pigs, cattle and zebrafish. Our studies revealed that nuclear localization of NPM2 was severely disrupted in fertilized eggs and 2cell embryos of KPNA2 KO females, while no effect was found in immature GV oocytes (Fig. 8B,C).

## MATERIAL and METHODS

### Generation of KPNA2 KO mice

For the generation of KPNA2 KO mice, ES cells with a gene trap mutation in the KPNA2 gene (clone XS0061) were purchased from MMRRC at University of California, Davis, and directly used for blastocyst injection. Germline chimeras were bred with C57Bl/6 mice and the colonies were maintained by breeding the resulting heterozygous mice. For genotyping of KPNA2 mice, a three primer PCR was developed. The sequences and conditions are listed in Table S2.

### Superovulation and embryo culture

Superovulation and embryo culture was performed as described elsewhere (Popova et al., 2005). Briefly, female mice were injected i.p. with 7.5 IU Pregmagon (Ceva) followed by injection of 7.5 IU Ovogest within 48-50 h. Mice were mated with C57Bl/6 males and checked for the presence of a copulatory plug on the following morning. Animals were sacrificed and oviducts were transferred to M2 medium (Sigma) containing hyaluronidase (0.1% w/v; Sigma). After release from the oviduct oocytes and embryos were cultured *in vitro* in M16 medium (Sigma) at 37°C under 5% CO_2_. For retrieval of GV oocytes, mice were injected with 7.5 IU Pregmagon and sacrificed 46 h after the injection. Follicles were retrieved from the ovary by punction with a needle, cumulus oocyte complexes (COC) were released and cumulus cells were removed by medium strongly vortexing for 5 min. For In vitro maturation of oocytes, COCs were isolated from the ovary and washed in IVM medium (TCM 199, 10% FCS, 0.23 mM sodium pyruvate, 50 IU PMSG). 20-30 COCs were transferred into drops with IVM medium covered with mineral oil and incubated for 14-15 h at 37°C under 5% CO2. For assessment of spontaneous oocyte activation, superovulated oocytes from not mated female mice were cultured in M16 medium and checked under the stereomicroscope or by DAPI staining for the formation of pronuclei.

### SDS PAGE and western blot

For western blot of mouse embryos, protein isolates of 80 oocytes or zygotes were loaded on a 10% SDS gel. For analysis of mouse tissues, 30 μg of tissue protein extracts were loaded on a 10% SDS gel. After transfer of proteins, the PVDF membrane was blocked by Odyssey blocking solution (LiCor, Bad Homburg, Germany) and subsequently incubated with primary antibodies at 4°C overnight. On the next day, the membrane was incubated with an IRDye-coupled secondary antibody for 1 h at room temperature and detection was performed using the Odyssey Infrared Scanner (LiCor, Bad Homburg, Germany). The generation of C-terminal and N-terminal antibodies against importin α7 was accomplished using standard protocols and has been described previously (Kohler et al., 1999). The generation of the KPNA6 antibody and the KPNA3 antibody is described elsewhere (Rother et al., 2011) The complete list of antibodies and conditions can be found in supplementary data (Table S3).

### RNA isolation and RT-PCR

Total RNA was isolated from mouse tissues or mouse preimplantation embryos using the TRIZOL method. Briefly, the material was homogenized in 1 ml Trizol reagent, 0.2 ml chloroform was added and after centrifugation, the colorless upper phase was transferred to a clean tube. For mouse tissues, 0.5 ml isopropyl alcohol was added, while for preimplantation embryos 0.5 ml isopropyl alcohol and 20 μg glycogen was added. After careful washing with ethanol, the RNA was air dried and dissolved in RNAse free water. The synthesis of cDNA was achieved by incubation in a 20 µl reaction mixture containing 200 U of MMLV Reverse Transcriptase (Promega) and 500 ng of random primer (Roche) at 25°C for 15 minutes, followed by 60 minutes incubation at 37°C and inactivation for 15 minutes at 70°C. The cDNA was diluted to 5 ng/μl. For PCR, a 25 µl reaction mixture consisted of 10 ng of the cDNA solution, 50 ng of each primer, 5 µM dNTP, and 1 IU Taq DNA polymerase (NEB). For detailed information on primer sequences and PCR conditions see Supplementary Information Table S2.

### β-Galactosidase staining of embryos

Cultured oocytes and embryos were rinsed in 0.1 M phosphate buffer pH 7.3 at room temperature, fixed for 5 minutes in fixation solution containing glutaraldehyde, washed 3 times for 5 minutes in wash buffer (phosphate buffer containing 1 mM MgCl2 and 0.02% NP40) and incubated with X-gal stain overnight at 37°C (for detailed protocol see http://www.med.umich.edu/tamc/laczstain.html). After staining, embryos were placed in wash buffer und stored at 4°C. For visualization, a Keyence microscope (Keyence, Bioreva BZ-9000, Germany) was used. Embryos of WT females mated to WT males were used for negative control.

### Immunofluorescence analysis of oocytes and embryos

Oocytes and preimplantation embryos of superovulated mice were collected, washed in PBS and fixed in 4% PFA in PBS at room temperature for 15 minutes. After permeabilization in PBS / 0.1% PVP (Sigma P0930) / 0.3% Triton X100 for 30 minutes at room temperature, oocytes and embryos were transferred to blocking buffer (containing PBS / 0.1% TX100 / 5% normal donkey serum) and incubated for 1h. For visualization of the meiotic spindle, oocytes were incubated with a FITC conjugated anti-α-tubulin antibody (Sigma; 1:100 in PBS / 0.1% PVP) for 2 h at 37°C. At the end of the incubation period, oocytes were washed in PBS / 0.1% PVP, and DNA was counterstained with 5 µg/ml Hoechst 33258 for 15 minutes and mounted with Fluorescence Mounting Medium. For immunofluorescence staining of all other proteins, the oocytes and embryos were incubated with the respective antibodies at 4°C overnight. After washing, embryos were incubated with secondary antibodies for 2 h, washed and DNA was counterstained using 5 µg / ml Hoechst 33258 for 15min. Finally, embryos were mounted with Fluorescence Mounting Medium and visualized under a confocal fluorescence microscope (Leica TCS SPE). Details on antibodies and conditions are listed in Supporting Information Table S3.

### Quantification of N/C signal ratio in stained embryos

For the quantification of the nuclear and cytoplasmic localization, representative pictures of stained embryos were analyzed using ImageJ software. Briefly, signal intensity in the nucleus and in the cytoplasm were measured and the ratio was calculated after subtraction of background signals.

### GST-importin pull-down assay

GST-importin pull-down assays were carried out as described earlier (Depping et al., 2008). GST and GST-fusion proteins were purified to almost homogeneity. GST protein served consistently as negative control. Briefly, GST or GST-importins were allowed to bind to glutathione-Sepharose 4B (GE Healthcare). In a typical experiment 100 μl beads were pre-equilibrated in IP-buffer (20 mM Hepes pH 7.5, 100 mM KOAC, 0.5 mM EGTA, 5 mM MgOAc, 250 mM sucrose, 4°C), mixed with 15 µg GST-fusion proteins followed by incubated at 4°C overnight. Oct4 was transcribed and translated *in vitro* in the presence of ^35^S-methionine (TNT Coupled Reticulocyte Lysate System, Promega, USA) according to the manufactureŕs protocol. After incubation, 10 µl of the reaction batch and His-tagged importin-β, were allowed to bind to the immobilised fusion-proteins. In competition experiments purified nucleoplasmin and importins were added in a 1:1 ratio. ^35^S-labelled methionine was obtained from Hartmann Analytic (Braunschweig, Germany). After washing sepharose beads were dissolved in 30 µl Laemmli buffer. Proteins were separated by SDS-PAGE (4-12%) and visualised by Coomassie Brilliant Blue staining. To detect the [^35^S]-labelled proteins, the dried gels were autoradiographed (16 h-24 h).

### Statistical analyses

Statistical analyses were performed with Prism 7 (Graphpad). Results are presented as means ± s.e.m. (body weight, quantification of immunofluorescence) or as means (litter size, embryonic development). In analyses comparing 2 groups, the unpaired two-tailed Student’s t-test was used for assessment of significance. In analyses comparing more than 2 groups, ANOVA was used to determine significance. For multiple comparisons, Dunnett’s multiple comparisons test was applied. A p-value < 0.05 was regarded as significant (*, p < 0.05; **, p < 0.01; ***, p < 0.001; ****, p < 0.0001; n.s., not significant).

## DISCUSSION

The concerted development of sperm and oocyte into a zygote marks the point, where two highest specialized cells form a new totipotent structure giving rise to every cell of the developing organism. This transition has to meet a variety of special needs regarding the integration of maternal and paternal chromatin, new synthesis of mRNA and proteins, establishment of cellular pathways and growth. Many of these processes require a coordinated nuclear transport, whose regulation is dependent on the expression of KPNAs, as they have the ability to bind different sets of cargoes. The systematic assessment of KPNAs during oocyte development has revealed their differential expression and suggested distinct functions during oogenesis (Mihalas et al., 2015). We have extended these analyses on the protein level showing that KPNA2, KPNA3 and KPNA6 are highly expressed in mouse oocytes as well as zygotes, while KPNA1 and KPNA4 show no expression. Our previous knockout mouse studies have revealed that maternal KPNA6 is crucial for early preimplantation development, while KPNA3 is completely dispensable at this developmental stage (Rother et al., 2011). To understand the role of KPNA2, we created KPNA2 KO mice. These mice display a severe phenotype of early embryonic developmental defect which confirms a study recently published by Wang et al. (Wang et al., 2023). In their study, Wang and colleagues observed a developmental 2cell stage arrest of embryos derived from KPNA2 KO females and suggested that the entry of the protein RSL1D1 into the pronuclei was impaired and the reason for the developmental arrest. Interestingly, in our investigation we noticed that the pronuclear localization of RSL1D1 was highly variable in KPNA2 KO zygotes, suggesting that even if RSL1D1 nuclear import was not completely abolished at this stage, a retarded nuclear transport could be an underlying mechanism. However, extending our analyses to 2cell stage embryos, the stage at which the developmental arrest finally occurs, did not reveal any differences in nuclear localization of RSL1D1 between WT und KO embryos anymore. Thus, although we do not rule out that RSL1D1 contributes to the embryonic arrest, we conclude, that further factors could be involved in the KPNA2-dependent phenotype.

Of the four maternally expressed KPNA paralogues, KPNA3 is totally dispensable for early embryonic development. The maternal depletion of KPNA2 or KPNA6 results in a total preimplantation embryo arrest (albeit the phenotypes show slightly different phenotypes), and embryos from KPNA7 KO mothers survive preimplantation period at 50% (Hu et al., 2010; Rother et al., 2011). Interestingly, KPNA7, not KPNA2, is specifically expressed in the ovary of cattle, humans and mice and its expression is the most abundant in cattle and human oocytes compared to other KPNA paralogues, while in mouse oocytes KPNA2 shows a higher expression than KPNA7 (Hu et al., 2010; Kelley et al., 2010; Tejomurtula et al., 2009; Wang et al., 2023). Thus, it is possible that human KPNA7 is the biological equivalent of mouse KPNA2 regarding its function in preimplantation development. This hypothesis is supported by data from Wang et al. showing, that the phenotype of KPNA2 KO mice could be rescued by microinjection of human KPNA7, but not mouse KPNA7 (Wang et al., 2023). These previous studies suggest that the function of KPNA paralogues is generally most important for early embryonic development but also indicate a certain plasticity of specific functions of KPNA paralogues during evolution.

Analysis of KPNA2 promoter activity and expression during early embryonic development revealed, that maternal KPNA2 is highly expressed in GV oocytes. The differing results of β-galactosidase activity, representing the transcription from the KPNA2 promoter on one hand and KPNA2 localization in MII oocytes on the other hand could be explained by a degradation of the (artificially expressed) β-galactosidase, while KPNA2 protein is still expressed in MII oocytes. In zygotes, maternal KPNA2 is translated from stored maternal mRNAs, defining KPNA2 as a maternal effect protein. Furthermore, a de novo transcription of KPNA2 from the embryonic genome is found in 2cell embryos and this transcription is reduced in 2cell embryos derived from KPNA2 KO females. Transcription of KPNA2 at the 2cell embryonic stage has been found in an earlier microarray analysis (Hamatani et al., 2004). Thus, we conclude, that KPNA2 is a maternally and embryonically expressed protein with maternal KPNA2 directly influencing the transcription of embryonic KPNA2. Our study shows that maternal KPNA2 is essential for embryonic development, while embryonic KPNA2 is dispensable, as KPNA2 KO embryos from heterozygous breedings survive.

In GV oocytes and zygotes maternal KPNA2 strongly localizes to the nuclei. Beginning at the 2cell embryo stage, we found varying localization of KPNA2 within the cell cycle, with phases of localization at the nuclear membrane and temporally strong nuclear accumulation. Currently it is unclear, if the localization of KPNA2 at the nuclear membrane, which can be found at each developmental stage until blastocyst, represents a short period of concerted entry of KPNA2 into the nucleus or if KPNA2 exerts a distinct function at the nuclear membrane. Interestingly, in addition to its function as a nuclear import mediator, KPNA2 has been shown to be involved in mitotic spindle formation (Guo et al., 2019). The abnormalities of the MII spindle in oocytes could indicate a role for KPNA2 also during meiotic spindle formation. Recent studies suggested that KPNA2 is a critical regulator of the spindle assembly factor TPX2 during mitosis (Guo et al., 2021). In this model KPNA2 sequesters TPX2 in the cytosol and timed phosphorylation of KPNA2 releases TPX2 to facilitate mitotic spindle formation. TPX2 has also been found to regulate the formation of the meiotic spindle and its premature presence interferes with proper spindle assembly (Brunet et al., 2008). Thus, the observed malformation of meiotic spindles in KPNA2 KO oocytes could be explained with the absence of regular KPNA2 levels affecting the temporally critical availability of TPX2.

We have shown that KPNA2 is a critical regulator of ZGA, an event that marks the de novo transcription from the embryonic genome. In mice, a first transcriptional wave can be observed beginning in late zygotes, while the majority of ZGA genes is transcribed in mid-to-late 2cell embryos (Aoki, 2022). Interestingly, the first wave of ZGA meanwhile has been regarded as a global low-level expression of thousands of genes throughout the embryonic genome, while the second wave represents a selective expression pattern. While dozens of genes have been shown to affect ZGA, the exact trigger of transcription initiation is not known yet, making it difficult to link KPNA2 and ZGA on a molecular level (Aoki, 2022). Besides KPNA2, KPNA6 has also been shown to be critically for ZGA (Rother et al., 2011). We therefore hypothesize that specific factors relying on nuclear transport could be the trigger of embryonic transcription initiation. KPNA2 and KPNA6 belong to different subfamilies, sharing only 48% identity, yet, both of them are essential for preimplantation development. On the other hand, KPNA2 shares 46% identity with KPNA7, the former being the closest relative of the latter, but KPNA7 is dispensable for development of preimplantation embryos (Wang et al., 2023). We therefore do not favour the hypothesis that ZGA relies on the collective action of all maternally expressed KPNAs (KPNA2, KPNA3, KPNA6 and KPNA7). In contrast, our observations in KPNA2 KO-and KPNA6 KO-derived embryos suggest that the affected ZGA-promoting factors are different ones, being a specific cargo of the respective KPNA paralogue, and their absence in the nucleus therefore can explain the differing phenotypes in KPNA2 and KPNA6 KO derived embryos.

In search for KPNA2-dependent targets we analyzed a number of maternal effect factors, proteins that are maternally derived and indispensable for the transition of the oocyte into an embryo. The transcription factor Oct4 is such a protein and has been found to be a master activator of ZGA in zebrafish (Lee et al., 2013; Leichsenring et al., 2013). In mice, Oct4 is a key regulator of the first wave of ZGA and knockdown in mouse oocytes leads to developmental stop at 4cell stage (Foygel et al., 2008; Tan et al., 2013). Considering the fact that Oct4 has to enter the nucleus, that its nuclear localization sequence is essential for the nuclear import and that it has been found to bind to KPNA2 in Y2H screens, pulldown assays and Co-IP experiments, we hypothesized that impaired Oct4 nuclear import could be the underlying mechanism for the developmental arrest in KPNA2 KO derived embryos (Li et al., 2008; Pan et al., 2004). However, by immunohistochemistry we could not find any differences in Oct4 localization between WT and KO. Using KPNA binding assays we furthermore showed that Oct4 binds to KPNA 1,2,3,4, and 6, suggesting, that the remaining KPNA paralogues are sufficient to transport Oct4 into the nucleus in GV oocytes, zygotes and 2cell embryos. Our binding assays are contrary to studies by Yasuhara et al. which have shown that in nuclear import assays Oct4 is transported by KPNA1, 2, and 4 only (Yasuhara et al., 2007), but not by KPNA3 and KPNA6. In this scenario, as KPNA1 and KPNA4 are not expressed in oocytes, KPNA2 would be the only remaining factor transporting Oct4 into the nucleus and its deficiency should severely affect the nuclear import of Oct4, which is not the case. Alternatively, other KPNA-independent mechanisms could exist that ensure Oct4 translocation into the nuclei of preimplantation embryos.

The transcription factor Sall4 is highly expressed in oocytes and early embryos and acts in concert with Oct4 and Nanog to regulate a number of metabolism-and transport-related genes necessary for early development (Tan et al., 2013). Maternal deletion of Sall4 results in developmental arrest of GV oocytes with incomplete meiosis resumption due to epigenetic misregulation (Xu et al., 2017). Another interesting candidate, the transcriptional activator Brg-1 has been one of the first identified maternal effect genes in mice and has been shown to be involved in ZGA regulation by modifying the chromatin structure. Like embryos from KPNA2 KO females, embryos from Brg-1-depleted females display a 2cell arrest (Bultman et al., 2006). The germ cell specific protein Stella has been shown to be involved in the epigenetic chromatin reprogramming of the zygote and maternally expressed Stella is crucial for the preimplantation development (Bortvin et al., 2004; Nakamura et al., 2007; Payer et al., 2003). However, all these three maternal effect proteins showed a regular (pro-)nuclear localization in embryos from KPNA2 KO females compared to WT, suggesting that other factors are either involved in their nuclear transport or can compensate for the KPNA2 loss. Moreover, we show that the dimethylation of lysine 9 of core histone H3 (H3K9me2) at the maternal pronucleus, which has been defined recently to be crucial for preimplantation development, was found to be unaffected in zygotes derived from KPNA2 KO females, suggesting a regular nuclear import and function of histone methyltransferase G9a (Au Yeung et al., 2019).

In contrast to this, the nuclear import of NPM2 was markedly disturbed in embryos from KPNA2 KO females. Expression of NPM2 is restricted to the ovary and has been shown to be crucial for the organization of the nucleolus-like body in GV oocytes and the nucleoli in early embryos (De La Fuente et al., 2004; Inoue et al., 2011). In *Xenopus* oocytes, nucleoplasmin is one of the most abundant proteins and has been implicated in sperm decondensation, while in mouse oocytes, its significance for sperm decondensation is still unclear (Vitale et al., 2007). Zygotes of female NPM2 KO mice display a reduced ability to cleave into 2cell stages defining NPM2 as a maternal essential factor (Burns et al., 2003). Moreover, NPM2 possesses a bipartite nuclear localization signal, representing a potential binding site for KPNAs. Our analyses of NPM2 localization in oocytes and early embryos of KPNA2 KO females revealed a severe nuclear transport defect. The fact, that the phenotypes of KPNA2 KO and NPM2 KO differ, could be explained with a partial inhibition of NPM2 nuclear import: other remaining KPNA paralogues could to some extent but not fully compensate for the import defect. Indeed, immunocytochemistry for NPM2 revealed a reduced but not completely abolished localization of NPM2 in the nuclei of oocytes and early embryos. Moreover, while embryos of NPM2 KO females sometimes survive the preimplantation period, embryos of KPNA2 KO females display an even stronger phenotype with a stop at the 2cell stage. Therefore we conclude, that additional factors must be relevant for the development of the phenotype observed in KPNA2 KO.

Interestingly, the coordinated nuclear import events in preimplantation embryos seem not only to depend on the availability of import factors and on their specificities towards certain substrates: a recent study by Nguyen and colleagues has suggested, that the timing of the nuclear entry of a protein also strongly depends on its biochemical affinity to karyopherins (Nguyen et al., 2022). In their study, the authors used *Xenopus laevis* embryos and studied the timely appearance of proteins in the nucleus which strongly correlated with the onset of their nuclear functions. Subsequently, the authors developed a model of competitive binding of substrates to a limited amount of karyopherin which correctly predicted the order of protein import found in cell-free droplets of *Xenopus* egg lysates with artificial nuclei. If this new model from *Xenopus* can be transferred to the mouse, it suggests, that depletion of KPNA2 could not only affect the specific nuclear import of KPNA2-dependent factors, but could also affect factors that are under normal conditions independent of KPNA2, because a KPNA2 deficiency could result in a global change of binding affinities and factors normally transported by KPNA2 might now bind to another KPNA paralogue thus influencing the import capacity of the latter. This scenario would also explain, why the nuclear import of proteins such as RSL1D1 and NPM2 is not totally abolished in embryos deficient of KPNA2.

In conclusion, we identify maternal KPNA2 as an essential protein for mouse early embryonic development enabling the correct start of ZGA. With NPM2 our study now adds a second factor to the list of proteins contributing to the embryo arrest in KPNA2 KO, besides RSL1D1. We infer that the observed phenotype has a multifactorial cause; future investigations could identify even more candidates contributing to the preimplantation arrest in KPNA2 KO.

## ACKNOWLEDGEMENTS

We thank Vivien Rabke and Steve Bomberg for help with animal caretaking. We wish to thank Michalina Alicka, Madeleine Skorna-Nussbeck, Anne Hahmann and Vivien Latuske for their excellent technical assistance. Lastl, we thank the Advanced Light Microscopy technology platform of the MDC for technical support.

